# N-Orbit: towards a universal model and metric for comparing tissue microenvironments

**DOI:** 10.1101/2025.04.23.650192

**Authors:** Barbara Xiong, Yuxuan Hu, Kai Tan

## Abstract

Advances in spatial omics technologies have greatly expanded our ability to explore tissue microenvironments (TMEs) across development and disease. Yet a theoretical framework for modeling and comparing tissue architecture in diverse biological contexts remains lacking. We introduce, *N-Orbit*, a mathematical model that captures both cell type composition and spatial relationships within tissue cellular neighborhoods, encoding this information in a vector format for computationally efficient distance metric calculations. While not a neighborhood detection method itself, N-Orbit enhances insights gleaned from the neighborhood generated by the plethora of recently developed methods. We benchmarked the N-Orbit-based neighborhood distance metric on spatial omics datasets containing ground truth neighborhoods and patient clinical outcomes. We demonstrated that the N-Orbit outperforms cell type enrichment (CTE)-based metrics in discriminating neighborhood types, predicting clinical variables, and identifying homologous structures across species. Additionally, N-Orbit enhances model interpretability by allowing neighborhoods to be traced back to their enriched N-Orbit structures. N-Orbit holds significant potential for deepening our understanding of how TMEs remodel during development, disease, and evolution.

## INTRODUCTION

The rapid advancement of spatial omics technologies has revolutionized the study of tissue microenvironments (TMEs) at high resolution. Imaging-based technologies, such as Multiplex Error-Robust Fluorescence In Situ Hybridization (MERFISH) for RNAs and Co-Detection-by-InDEXing (CODEX) for proteins, use multiplexed *in situ* hybridization of fluorescent probes.^1,2^ Sequencing-based technologies, including STARmap and Stereo-Seq, employ microarrays of spatially barcoded probes to measure RNA expression with single-cell spatial resolution.^3,4^ By leveraging spatial transcriptomic and proteomic data, researchers can map the diverse landscape of intermingling cell types, uncovering spatial heterogeneity and cell-cell interactions. This approach holds great promise for uncovering valuable insights into both healthy development and disease progression.

Tissue cellular neighborhoods (TCNs) or spatial domains are fundamental organizational units of tissue architecture. They are spatially contiguous tissue regions with a consistent composition of cell types that perform coordinated function(s) through cell-cell interactions. The phenotype of a TCN reflects the collective behavior of its constituent cells, focusing on how they communicate and organize to perform specialized functions. Identifying TCNs is crucial for understanding tissue development, homeostasis, and disease mechanisms. Several computational tools have been developed to detect TCNs based on various spatial omics data, including our recently established method, CytoCommunity.^5–10^ Partitioning spatial omics maps into TCNs enables the identification of such localized patterns that could otherwise be diluted and overlooked when analyzing spatial relationships in bulk. This is particularly important in contexts with high spatial heterogeneity. The study of TCNs reveals how spatial heterogeneity is organized, which is often hierarchical. For example, the splenic B cell zone consists of germinal centers polarized into light and dark zones, composed of centrocytes and centroblasts, respectively.

Analyzing TCNs over time may capture dynamic processes such as cellular proliferation, migration, and reorganization, all of which are crucial to understanding disease progression. As diseases develop and progress, TCNs may undergo significant alterations. Monitoring these changes provides insights into disease pathophysiology and help evaluate and predict treatment efficacy and clinical outcomes. For example, Schürch, *et al*. found that enrichment of PD-1+CD4+ T cells within a granulocyte-associated TCN correlated with improved survival in high-risk colorectal cancer patients.^11^ Similarly, Reiss, *et al*. identified CD68+CD163+ macrophage-enriched TCNs that were associated with poor clinical outcomes in patients with diffuse large B-cell lymphoma (DLBCL).^12^

The comparison of TCNs between species can also provide critical insights into the conserved roles and behaviors of cellular organizations across evolutionary time. While individual cell types may vary slightly between species in terms of structure or function, analysis of conserved TCNs allows us to better understand the evolutionary selective pressure on cellular communication and conservation of functional tissue architecture. For instance, comparing tertiary lymphoid organs between human and mouse may help identify certain cellular interactions that are crucial for the effective functioning of this type of TCNs.

While the problem of TCN detection itself has been thoroughly explored, a fundamental open question is how to quantitatively characterize the changes in resulting TCNs across various conditions (e.g. treatment periods, cancer risk subtypes), a challenge that holds similar significance and broad applicability to differential gene expression (DGE) analysis. Developing quantification methods for application on spatially or temporally discrete samples is particularly difficult, as these samples are often derived from physically distinct tissues. The absence of physical continuity and the limitation to a few clinically relevant time points make it difficult to trace spatial rearrangements. As a result, previous attempts have primarily analyzed TCNs only at the level of cell-type enrichment. There has been a lack of principled mathematical models for describing TCN structure. It is infeasible to directly align individual cells from one sample to another to track spatial changes. Furthermore, no methods exist to compare TCNs across species, a challenge analogous to the multiple sequence alignment (MSA) problem for homology detection.^13^ Alignment-based methods are limited by their computational complexity and sensitivity to highly divergent sequences. However, *k-mer*-based methods, which consider word frequencies, have proven to be an effective heuristic technique with substantially better robustness and scalability.^14^ We draw inspiration from this approach and adapt it to the concept of TCNs.

Here, we introduce the *N-Orbit*, an interpretable mathematical model for the building block of TCNs. We draw an analogy to atomic physics, where cells correspond to subatomic particles, N-Orbits to atoms, and TCNs to molecules. The N-Orbit model is applicable to single-cell resolution spatial omics data, where each cell is assigned a cell type and its spatial coordinates. By applying the N-Orbit model, we define a distance metric to quantify differences between pairs of TCNs in an alignment-free manner. We optimize the N-Orbit structure to best capture essential spatial relationships between cell types, providing a foundation for computationally efficient neighborhood distance calculation. Using the neighborhood distance metric, we can construct an overall distance matrix for whole datasets, allowing for the projection of neighborhoods onto a common low-dimensional embedding for visualization and global comparison. We also formulate an N-Orbit enrichment test, allowing for the traceback of TCN changes between conditions to the relevant N-Orbit structures for enhanced interpretability. Just as DGE analysis is used to identify distinct cell phenotypes, our N-Orbit approach can identify TCN-level phenotypes.

We systematically evaluated the N-Orbit model and its associated distance metric through several key assessments. First, we tested its ability to effectively distinguish analogous neighborhoods from unrelated ones. Second, we explored the robustness of the model across a range of hyperparameters and neighborhood detection methods. Third, we examined its utility in capturing biologically and clinically relevant information. Overall, we demonstrated the potential of this framework to augment the insights gained from the myriad of recently developed neighborhood detection methods, as well as its broad applicability across diverse TME contexts.

## RESULTS

### Overview of the N-Orbit framework

We derived N-Orbit structures from n-hop subgraphs of labeled k-Nearest Neighbors graphs based on spatial coordinates and annotated types of each cell (Figure 1A). Unlike traditional n-hops, which are induced subgraphs of nodes within *n* edges of a target node, N-Orbits encode only the sets of cell types within *N* levels of proximity to each cell, without capturing the exact subgraph connections. This construction allows N-Orbits to be easily enumerated, reduces the number of possible combinations (thus increasing repetitions of structures), and makes them representable as vectors for computational analysis. Each vector encodes both the identity of the center cell type (the “*nucleus”*) and the proximity level of each neighboring cell type. With this vector formulation, the distance between two N-Orbits can be computed algebraically by their Manhattan distance (Figure 1B). The distance between two neighborhoods is then defined as the minimum total distance between the set of N-Orbits representing each neighborhood (Figure 1C). These pairwise neighborhood distances can be compiled into an overall neighborhood distance matrix (Figure 1D) and sample-level distances can be calculated analogously (see Methods).

**Figure 1:**
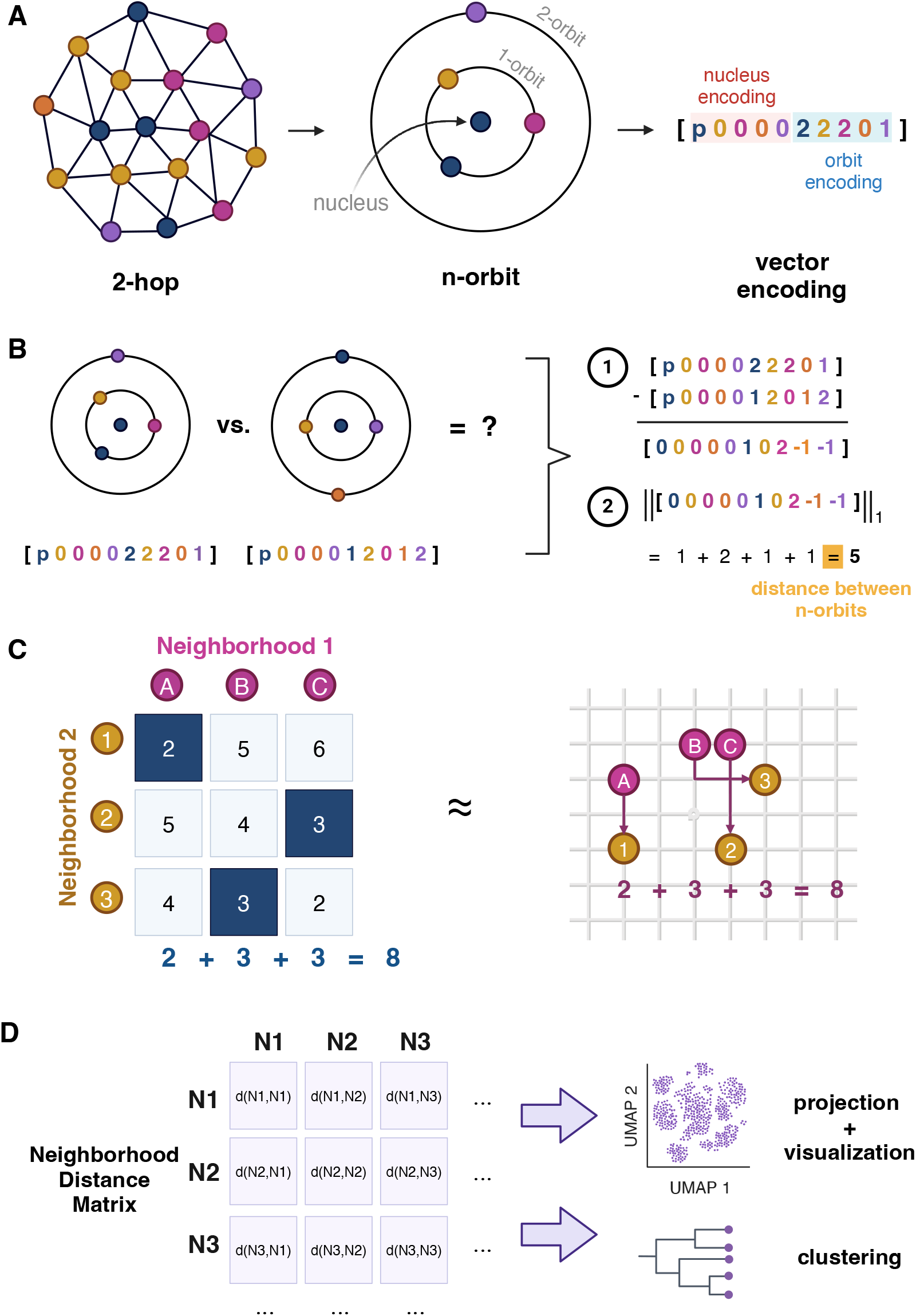
Schematic of the N-Orbit framework. **A)** Derivation of N-Orbits from 2-hop subgraphs of spatial k-Nearest Neighbors graphs labeled by cell-type, along with corresponding vector encoding. The vector encoding comprises nucleus and orbit components where each index corresponds to a possible cell type. The nucleus penalty change *p* is applied at the nucleus encoding index corresponding to the center cell type, with all other elements set to zero. Cell types in the 1- and 2-orbit are denoted with 2 and 1 in the orbit encoding, respectively, while cell types not included are denoted as zero. **B)** Example calculation of distance between two N-Orbit structures as the Manhattan distance between their vector representations. **C**) Example calculation of N-Orbit-based distance between two neighborhoods as the minimum total distance between the N-Orbits in neighborhood 1 and neighborhood 2, determined by a cost matrix optimization. **D**) Compilation of the N-Orbit-based neighborhood distance matrix, which serves as an input for projection, visualization, and clustering analysis. Created with BioRender.

We compared the performance of our N-Orbit-based neighborhood distance metric to that of a naïve approach based on cell type enrichment (CTE) (see Methods). We assessed the ability of each neighborhood distance metric to discriminate between pairs of neighborhoods of varying relatedness with two-sample t-tests and the Area Under the Receiver Operating Characteristic (AUROC) of distance-thresholded classifiers. Due to the oversensitivity of t-tests to large sample sizes (i.e. neighborhood pairs), and to more accurately quantify statistical significance between the performance of N-Orbit and CTE, we focused on AUROCs and corresponding DeLong test p-values as our primary performance metrics. P-values of t-tests are provided for reference in Supplementary Table 2.

### Evaluation of the N-orbit model using synthetic neighborhoods

We first simulated synthetic neighborhoods to validate the N-Orbit model’s ability to capture spatial cell type relationships beyond global cell type composition (Figure 2A-E). We designed synthetic neighborhood types consisting of 3 cell types (CT1, CT2, CT3) with similar marginal distributions (reflecting overall cell type composition) but different pairwise joint distributions (representing cell type interactions). Three broad classes of neighborhoods (A, B, and C) were generated using a Poisson process model (Methods), each with varying marginal distributions. Within each class, one neighborhood subtype (A1, B1, C1) had independently distributed cell types, while another subtype (A2, B2, C2) had a mixture of monotypic or heterotypic cell type interactions. These neighborhood subtypes were then grouped into two sample types (X, Y), each containing one neighborhood type from each broad class. These sample types were designed to model clinical variability that may influence neighborhood structure in real patient datasets.

**Figure 2:**
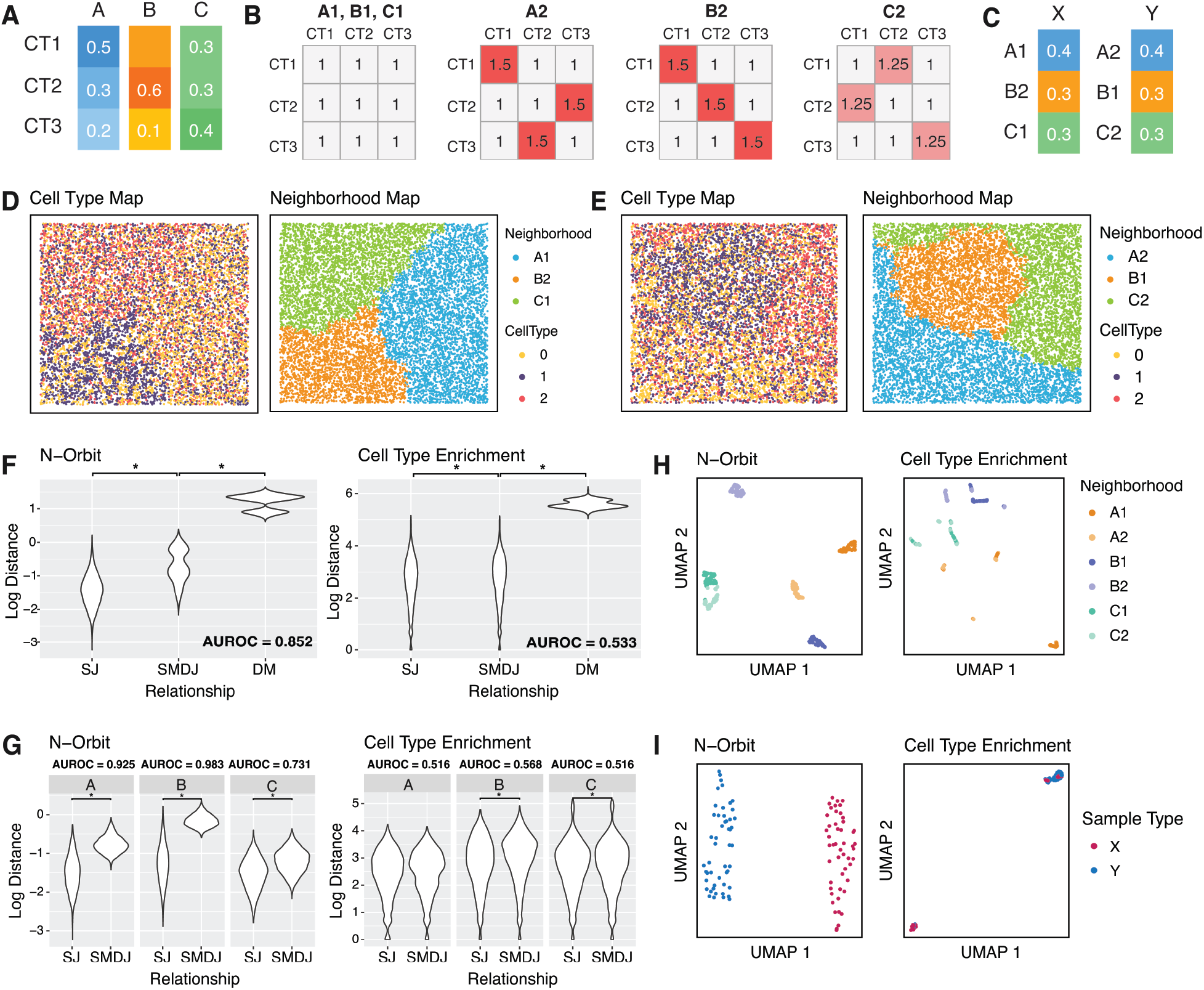
Performance evaluation of N-Orbit model to distinguish synthetic neighborhoods of similar marginal but different joint distributions. **A)** Cell type (CT) compositions for each neighborhood class A, B, and C. **B)** Cell type interactions (edge potentials) for each neighborhood. **C)** Proportions of each neighborhood within sample types X and Y. **D)** Example cell type and neighborhood maps for Sample Type X. **E)** Example cell type and neighborhood maps for Sample Type Y. **F)** Violin plots of log (scaled) N-Orbit and Cell Type Enrichment (CTE)-based neighborhood distances for neighborhood pairs with same joint (SJ), same marginal but different joint (SMDJ), and different marginal (DM) distributions. Asterisks indicate statistical significance by t-test. The area under the receiver operating characteristic curve (AUROC) are provided for SJ vs. SMDJ comparison. **G)** Violin plots of log (scaled) N-Orbit and CTE-based neighborhood distances for SJ and SMDJ neighborhood pairs within each neighborhood class. Asterisks indicate statistical significance by t-test. AUROC are provided for SJ vs. SMDJ comparison. **H)** UMAPs of N-Orbit and CTE neighborhood distance matrices, color-coded by neighborhood. Each point represents one neighborhood. **I)** UMAPs of N-Orbit and CTE sample distance matrices, color-coded by sample type. Each point represents one sample.

When aggregating all pairwise neighborhood distances across all neighborhood classes, the N-Orbit distance metric significantly outperformed the CTE distance in discriminating between neighborhood pairs of the same joint distribution (SJ) from pairs of the same marginal but different joint distributions (SMDJ) (AUROC: N-Orbit = 0.85, CTE = 0.53; DeLong test p-value < 1e-324) (Figure 2F-G). Within each broad neighborhood class, there was also a substantially greater overlap in neighborhood distances between SJ and SMDJ pairs using the CTE distance (AUROC: overall = 0.53, A = 0.52, B = 0.57, C = 0.52) compared to the N-Orbit distance (AUROC: overall = 0.85, A = 0.93, B = 0.98, C = 0.73). Performance of the N-Orbit distance for this task was significantly superior to the CTE distance (DeLong test p-value: A < 1e-324, B < 1e-324, C = 1.37e-86). Uniform Manifold Approximation and Projections^15^ (UMAPs) based on neighborhood distance metrics revealed clear separation of neighborhood subtypes within the same broad class using the N-Orbit distance metric but not the CTE distance metric (Figure 2H). While separation between C1 versus C2 neighborhoods was weaker in the N-Orbit-derived UMAP, it remained discernible, reflecting the design of C2 with weaker cell type interactions than A2 and B2. Additionally, sample-level UMAPs exhibited perfect separation between sample types X and Y using the N-Orbit distance metric, while the CTE distance metric resulted in intermixing (Figure 2I). Overall, these results underscore the N-Orbit distance metric’s superiority in capturing subtle nuances in cell type interactions compared to the naïve CTE distance metric.

We simulated another dataset that contained 8 cell types and 5 neighborhood types, each with a uniform cell type distribution but varying pairwise joint distributions (Supplementary Figure 1A-C). Neighborhood type N1 featured independently distributed cell types, while N2-N5 had various cell type interactions. These neighborhood types were grouped into two sample types, X (comprising N1, N2, and N4) and Y (containing N1, N3, and N5). Again, the N-Orbit distance metric significantly outperformed the CTE distance in distinguishing neighborhood pairs of the same type from those of different types (AUROC: N-Orbit = 0.80, CTE = 0.52; p-value < 1e-324) (Supplementary Figure 1E). Principal component analysis (PCA) plots at both the neighborhood and sample levels also demonstrated much better separation of neighborhood and sample types when using N-Orbit distance compared to the CTE distance (Supplementary Figure 1F-G).

### Evaluation of N-Orbit model on manually annotated mouse TCNs

#### Mouse spleen TCNs

Goltsev *et al*. provide a CODEX dataset of 3 healthy mouse spleens, manually annotated into four anatomical TCN types: red pulp, B-zone, marginal zone, and peri-arterial lymphoid sheath (PALS) (Figure 3A).^16^ We used these manually annotated neighborhoods as ground truth labels to assess whether our N-Orbit distance metric could accurately discriminate neighborhoods of the same type (across CODEX images) from neighborhoods of different types (Figure 3B). Both the N-Orbit and CTE distance metrics perfectly separated neighborhood pairs of the same type from those of different types. To increase our effective sample size and account for batch effects, we split each neighborhood type within each sample into separate neighborhood *instances* based on connected components of the k-Nearest Neighbors spatial graph, yielding 90 total instances across the dataset. We then compared the distances between neighborhoods of similar and dissimilar types within the same sample (Figure 3C). The N-Orbit-based distance metric significantly outperformed the CTE metric (AUROC: N-Orbit = 0.92, CTE = 0.70; p-value = 2.61e-50). Additionally, projecting each neighborhood distance matrix via UMAP demonstrated clearer separation between neighborhood types using the N-Orbit distance than the CTE distance (Figure 3D). The performance of the N-Orbit distance, as measured by AUROC, was also stable across hyperparameters, peaking at *N* = 1, *p* = 25, and *s* ≥ 1000 (Supplementary Figure 2A-B).

**Figure 3:**
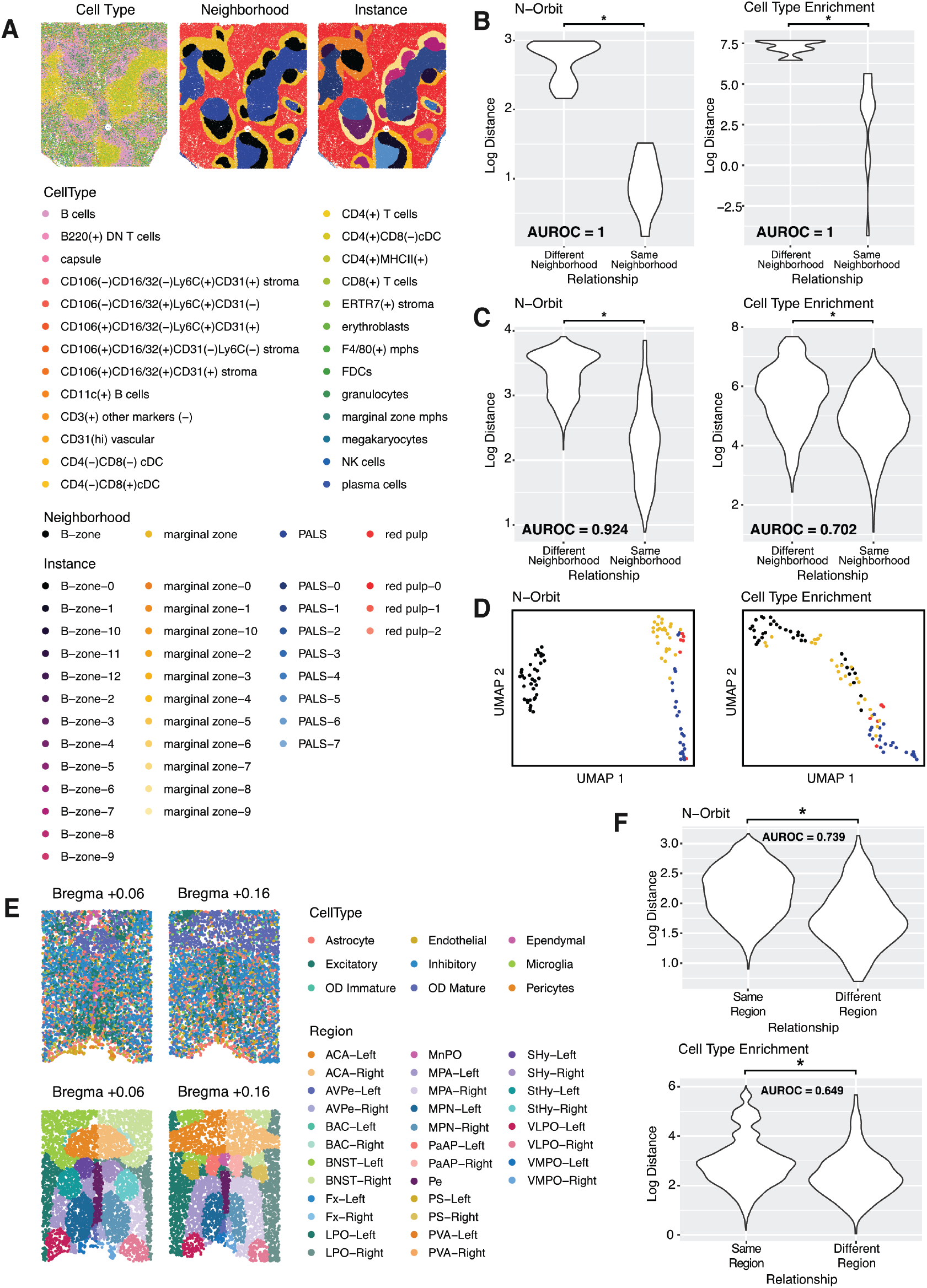
Performance evaluation of N-Orbit model to distinguish mouse anatomical regions. **A)** Cell type, neighborhood, and neighborhood instance maps for one mouse spleen CODEX sample. **B)** Violin plot of log (scaled) N-Orbit and CTE-based neighborhood distances between neighborhood pairs of different vs. similar types, with corresponding AUROC values. **C)** Violin plot of log (scaled) N-Orbit and CTE-based neighborhood distances between neighborhood instances of different versus similar types, with corresponding AUROC values. **D)** UMAPs of N-Orbit and CTE-based neighborhood instance distance matrices, color-coded by neighborhood type. **E)** Cell type and region maps for two mouse hypothalamus MERFISH samples. **F)** Violin plots of log (scaled) N-Orbit and CTE-based neighborhood distances between regions of similar vs. different mouse hypothalamic nuclei, with corresponding AUROC values.

We also assessed the robustness of the N-Orbit distance metric to upstream neighborhood detection methods previously benchmarked by Hu *et al*. (Supplementary Figure 2C).^6–10^ In Hu *et al*., CytoCommunity was the most accurate in generating neighborhoods that closely matched the ground truth, particularly in identifying the marginal zone. Other methods failed to detect a spatially coherent marginal zone, with stLearn partitioning cells only into the other three neighborhood types. Both CytoCommunity and UTAG (the second-best performing method, which adequately captured the other three neighborhood types) achieved near-perfect AUROC values in discriminating neighborhoods of similar and different types across samples when using the N-Orbit distance metric (AUROC= 1.0 and 0.96, respectively). Other neighborhood detection methods also performed reasonably well (AUROC BayesSpace = 0.85, STAGATE = 0.79, stLearn = 1.0, SpatialLDA: 0.668). Lastly, we applied the N-Orbit distance metric to compare neighborhoods generated by each neighborhood detection method to the corresponding ground truth neighborhoods (Supplementary Figure 2D). Consistent with the findings from Hu *et al*., CytoCommunity-generated neighborhoods exhibiting the closest proximity to the ground truth.

#### Mouse hypothalamus TCNs

We performed a similar analysis on manually annotated MERFISH mouse hypothalamic nuclei from Moffitt et al.^17^ This dataset comprises 17 neighborhood types across 5 mouse hypothalamus slices. To take advantage of the bilaterally symmetric nature of the anatomy, we divided non-midline structures into left and right neighborhood instances, resulting in a total of 101 instances across the dataset (Figure 3E). We then evaluated the ability of the N-Orbit and CTE distance metrics to distinguish neighborhood instances derived from the same versus different hypothalamic nuclei (Figure 3F). The N-Orbit distance metric significantly outperformed the CTE metric in discerning between similar and dissimilar pairs of neighborhood instances (AUROC: N-Orbit = 0.739, CTE = 0.649; p-value = 5.30e-9). In addition, the performance of the N-Orbit distance metric, as measured by AUROC, remained stable across hyperparameters, peaking at *N* = 1, *p* = 5, and *s* ≥ 1000 (Supplementary Figure 2E-F).

#### Mouse colitis TCNs

We also assessed the ability of the N-Orbit distance metric to distinguish intestinal layer-based neighborhoods in a mouse colitis model from Cadinu *et al*. (Figure 4A-B).^18^ This dataset comprises 52 MERFISH samples collected from 15 mice across four time points (days 3, 6, 9, and 21) after induction of colitis with dextran sulfate sodium (DSS). The authors provided neighborhood labels based on intestinal layers–lumen [LUM], mucosa [MU], follicles [FOL], submucosa [SM], muscularis externa [ME], and mesentery [MES]–as well as their relative order of emergence, designated numerically. This labeling scheme provided a hierarchical organization to the neighborhood types by intestinal layer (e.g. MU vs. SM), followed by specific neighborhood labels (e.g. MU1 vs. MU2). Across the dataset, there were 685 individual neighborhoods in total. We tested the N-Orbit distance metric’s performance at both the intestinal layer and neighborhood label levels (Figure 4C-D). The N-Orbit distance metric significantly outperformed the CTE distance metric in distinguishing neighborhood pairs from the same versus different intestinal layers (AUROC: N-Orbit = 0.740, CTE = 0.54; p-value < 1e-324). Additionally, N-Orbit demonstrated superior performance in distinguishing neighborhoods with the same label (AUROC: N-Orbit = 0.90, CTE = 0.70; p-value < 1e-324). Furthermore, the N-Orbit-derived UMAP revealed better separation of intestinal layers and neighborhood labels compared to those derived from CTE distances (Figure 4E), particularly highlighting the highest diversity within the mucosal neighborhoods. The UMAP generated from the sample-level N-Orbit-based neighborhood distance matrix also showed clear separation of samples by time post-DSS colitis induction (Figure 4F).

**Figure 4:**
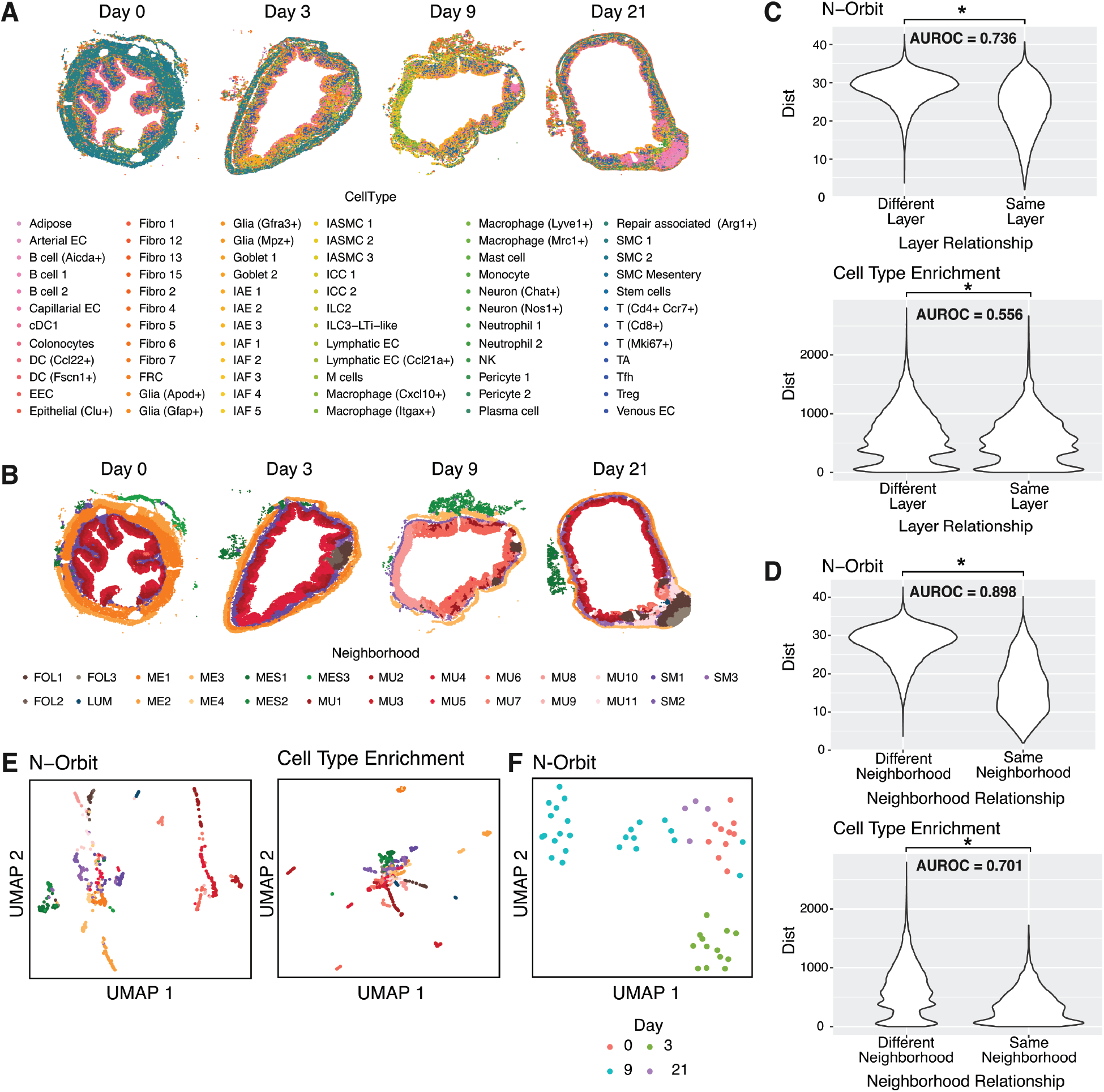
Performance evaluation of N-Orbit model to distinguish intestinal layers and neighborhoods from mouse colitis MERFISH data. **A)** Cell type maps of one MERFISH sample from each day post dextran sulfate sodium (DSS)-induced colitis. **B)** Neighborhood maps of one MERFISH sample from each day post DSS-induction of colitis. **C)** Violin plots of (scaled) N-Orbit and CTE-based neighborhood distances between neighborhood pairs of the same versus different intestinal layers, with corresponding AUROC values. **D)** Violin plots of (scaled) N-Orbit and CTE-based neighborhood distances between neighborhood pairs of the same versus different neighborhood labels, with corresponding AUROC values. **E)** UMAPs of N-Orbit and CTE-based neighborhood distance matrices, color-coded by neighborhood label. Each point represents a neighborhood. **F**) UMAP of N-Orbit-based sample distance matrix, color-coded by day post DSS-induction of colitis. Each point represents a sample.

### Therapy response prediction using N-Orbit distances

#### TNBC chemotherapy with or without immunotherapy

We evaluated the utility of the N-Orbit distance metric in predicting treatment outcomes using two triple-negative breast cancer (TNBC) datasets. The first dataset, from Wang *et al*., comprises 1855 imaging mass cytometry samples from 279 TNBC patients who were treated with chemotherapy with or without anti-PD-L1 immunotherapy, at three time points (Baseline, On-Treatment, Post-Treatment).^19^ Treatment outcomes for each patient were classified as either pathological complete response (pCR, 150 patients) or residual disease (RD, 129 patients) (Figure 5A-B). Using CytoCommunity supervised by treatment phase as the condition label, we identified 14,718 individual neighborhoods across the dataset. The UMAP of the N-Orbit-based neighborhood distance matrix revealed distinct regions of density corresponding to neighborhoods from different treatment phases (Figure 5C, Supplementary Figure 3). To predict treatment outcomes based on individual neighborhoods, we trained a support vector machine (SVM) classifier using PCA-projected vectors (# components = 35) of the neighborhood distance matrix, treatment type, and treatment phase as inputs. Using the N-Orbit distance metric, this classifier achieved an average test accuracy of 76.7 +/-1.1% (15-fold cross validation), compared to 68.6 +/-1.2% for the CTE distance metric (Figure 5D). Given that individual neighborhoods alone are unlikely to predict treatment outcomes, we also evaluated test accuracy after ensembling predictions from all neighborhoods in a patient’s samples via majority vote. By ensembling predictions from neighborhoods across all treatment phases, the N-Orbit-based classifier achieved a test accuracy of 97.8%, significantly outperforming the CTE-based classifier which achieved an accuracy of 80.8%. Additionally, when ensembling predictions from Baseline neighborhoods only, the N-Orbit-based classifier maintained a high test accuracy of 98.6%, compared to 78.4% with the CTE distance. These results suggest that the N-Orbit distance metric can predict treatment response with very high accuracy even before the onset of treatment. Performance was consistent across different N-Orbit depths, with optimal results observed at *N* = 2 (Supplementary Figure 2G).

**Figure 5:**
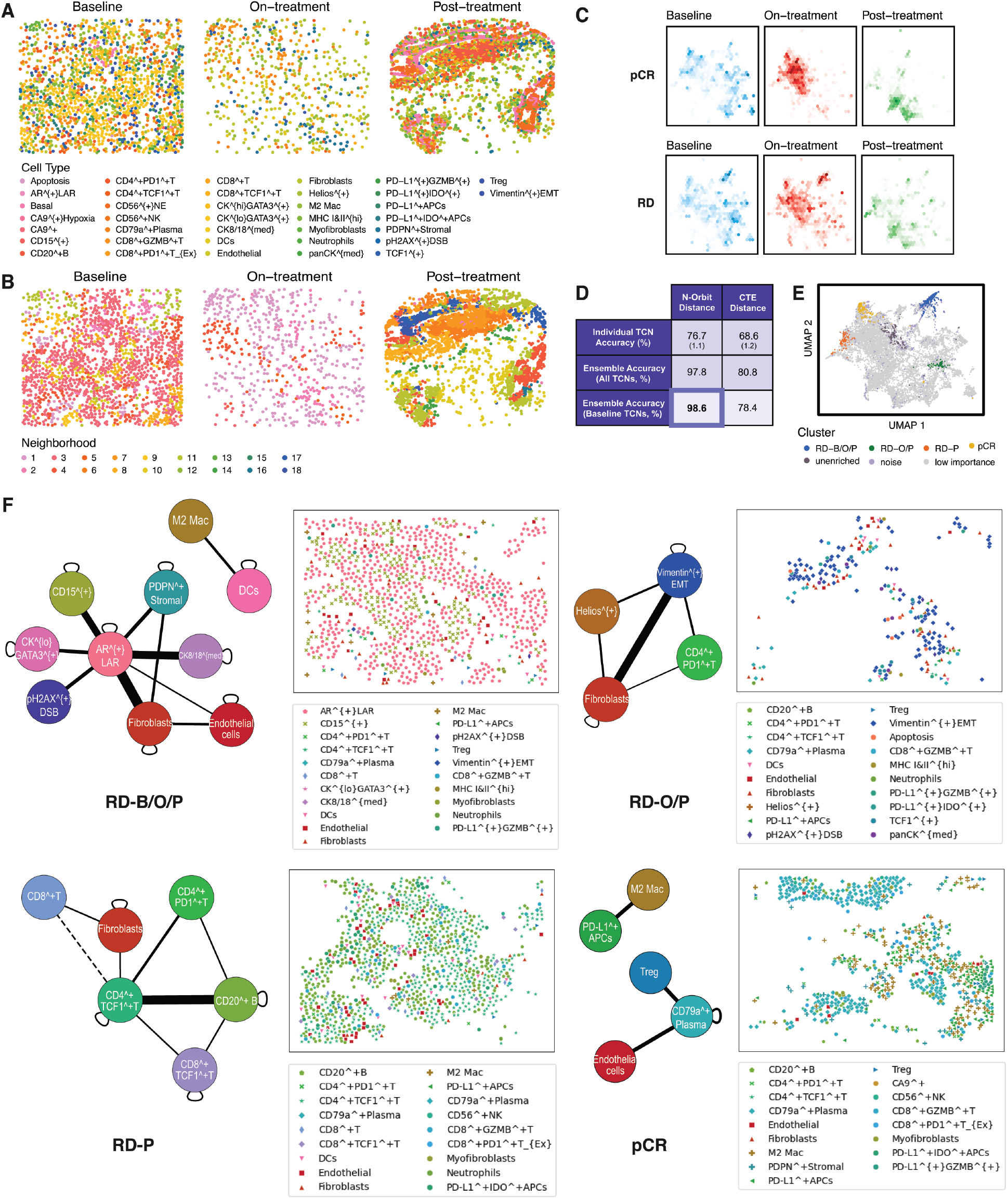
Application of N-Orbit distance to predict treatment response in triple-negative breast cancer (TNBC) patients treated with chemotherapy with or without immunotherapy. **A)** Cell type maps for samples from each treatment phase (Baseline [B], On-treatment [O], Post-treatment [P]) for one TNBC patient. **B)** Neighborhood maps for samples from each treatment phase for the same TNBC patient. **C)** UMAP density plots for the N-Orbit-based neighborhood distance matrix, faceted by treatment phases as columns and treatment outcomes as rows. pCR, pathological complete response; RD, residual disease. Density is computed from the number of neighborhoods within each hexagon, with colors scaled with respect to density ranges within each treatment phase. **D)** Performance statistics for treatment outcome predictions from N-Orbit and CTE-based classifiers. Standard deviations are shown in parentheses. **E)** Enriched neighborhood clusters (RD-B/O/P, RD-O/P, RD-P, pCR) plotted on the UMAP of the neighborhood distance matrix. Unenriched, noise (as determined by HDBSCAN), and low importance neighborhoods (as determined by loading contribution to RFE-selected PCA features) are plotted in gray for reference. **F)** Pruned summary graphs of enriched neighborhood clusters, derived from their enriched N-Orbits, with corresponding example cell type maps. Solid edges indicate 1-orbit relationships between N-Orbits, with edge weights indicating relative frequency of cell type interactions among enriched N-Orbits. Dashed edges indicate 2-orbit relationships. Self-edges indicate monotypic interactions of that cell type.

Feature selection using recursive feature elimination and PCA loading traceback identified the top 1,406 non-reduced features (equivalently, neighborhoods) relevant for predicting treatment outcomes from Baseline neighborhoods. Clustering of the distance matrix between these neighborhoods using HDBSCAN^20^ identified three RD-enriched and one pCR-enriched neighborhood clusters (Figure 5E). One RD-enriched cluster (RD-B/O/P, 433 neighborhoods) was enriched across all three treatment phases (p = 1.41e-40), while another (RD-O/P, 76 neighborhoods) was enriched particularly after treatment onset (p = 1.07e-5). The remaining cluster (RD-P, 99 neighborhoods) was enriched post-treatment (p=0.00867). In contrast, the pCR-enriched cluster (146 neighborhoods) was enriched primarily at Baseline and On-treatment time points (p=1.84e-6).

To enhance interpretability of clinically informative N-Orbit features, we developed a bootstrap-permutation test to identify N-Orbits enriched in each neighborhood cluster and compiled the results into a *pruned summary graph* (Figure 5F, Supplementary Figures 4-8). The subgraph for the RD-B/O/P cluster included 7 cell types centered around the AR^+^LAR malignant cell type, the most prevalent cell type in this cluster, alongside interactions between dendritic cells and M2 macrophages. Other cell types interacting with AR^+^LAR malignant cells included CD15^+^ malignant cells, as well as PDPN+ stromal cells, fibroblasts, and endothelial cells, which formed various cliques with the AR^+^LAR malignant cells. Notably, while Wang *et al*. previously identified the enrichment of CD15^+^ malignant cells in on-treatment resistant tumors, our N-Orbit analysis further highlights their interaction with AR^+^LAR malignant cells within resistance-enriched neighborhoods. The RD-O/P cluster demonstrated clique formation involving Vimentin+ epithelial-to-mesenchymal transition (EMT) malignant cells, fibroblasts, Helios+ cells and CD4+ PD-1+ T-cells. Cancer-associated fibroblasts (CAFs) have been implicated in driving vimentin expression and EMT in breast cancer, which is pertinent to metastasis.^21,22^ Helios expression also promotes the stability of regulatory T cells, which may counteract the effects of immunotherapy.^23^ The RD-P cluster was centered around CD4+ TCF1+ T cells interacting with various other lymphocyte types and fibroblasts. CAFs have been noted to suppress T cell function through extracellular matrix remodeling and promotion of immune checkpoints.^24,25^ In contrast, the pCR cluster included interactions between CD79a+ plasma cells, endothelial cells, and regulatory T cells, alongside interactions between M2 macrophages and PD-L1+ antigen-presenting cells (APCs). This differs from the plasma cell interactions with malignant epithelial cells noted in resistant tumors by Wang *et al*. and contrasts with the M2 macrophage interactions with dendritic cells, another type of APC lacking PD-L1 expression, in RD-B/O/P.

#### TNBC pembrolizumab plus radiation therapy

The second dataset, from Shiao *et al*. includes 78 CODEX samples from 28 TNBC patients treated with the immune checkpoint inhibitor (ICI) pembrolizumab (pembro) in combination with focal radiation therapy (RT) (Figure 6A-B).^26^ Samples were collected at three time points: before treatment (T1), after the first cycle of pembro (T2), and following the second cycle of pembro with RT (T3). Shiao *et al*. identified two responder subgroups (R1, 8 patients; R2, 13 patients) and a non-responder subgroup (NR, 7 patients). Using unsupervised CytoCommunity, we identified 629 neighborhoods across the dataset. Using a similar procedure to the previous TNBC dataset, we applied an SVM classifier to predict treatment response based on dimension-reduced neighborhood distance vectors and treatment phase (Figure 6C-D, Supplementary Figure 9). We also ensembled neighborhood-wise predictions to make overall treatment response predictions for each patient. Our N-Orbit-based classifier achieved a macro-F1 score of 0.691 +/-0.059 for predicting response type from individual neighborhoods, outperforming the CTE metric, which yielded a macro-F1 score of 0.505 +/-0.079. After ensembling predictions from all neighborhoods, the macro-F1 improved to 0.845 using N-Orbit distances (0.794 with CTE distances). Notably, when restricted to T1 neighborhoods only, the ensembled N-Orbit predictions maintained a high macro-F1 of 0.838 (compared to 0.611 with CTE distances). Performance was consistent across tested N-Orbit depths, with optimal results at both *N* = 2 and *N* = 3 (Supplementary Figure 2H).

**Figure 6:**
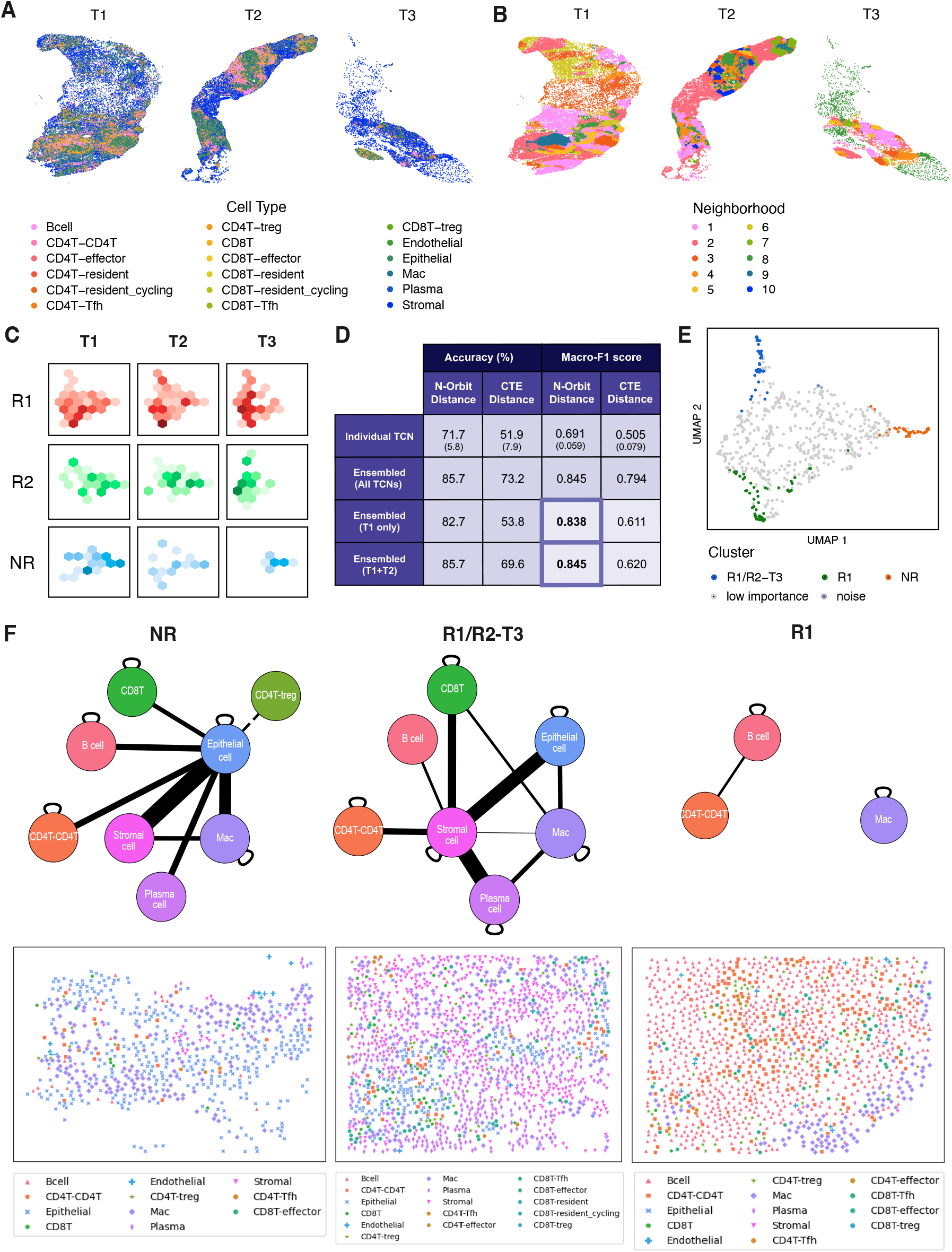
Application of N-Orbit distance to predict treatment response in triple-negative breast cancer patients (TNBC) treated with neoadjuvant pembrolizumab and radiation therapy. **A)** Cell type maps for samples from each treatment phase for one TNBC patient. **B)** Neighborhood maps for samples from each treatment phase for the same TNBC patient. **C)** UMAP density plots for the N-Orbit-based neighborhood distance matrix, faceted by treatment phases (T1, T2, T3) as columns and treatment outcomes (Response 1 [R1], Response 2 [R2], No Response [NR]) as rows. Density is computed from the number of neighborhoods within each hexagon, with colors scaled with respect to density ranges within each treatment outcome. **D)** Performance statistics for treatment outcome predictions from N-Orbit and CTE-based classifiers. Standard deviations are shown in parentheses. **E)** Enriched neighborhood clusters (NR, R1/R2-T3, R1) plotted on the UMAP of the neighborhood distance matrix. Noise (as determined by HDBSCAN) and low importance neighborhoods are plotted in gray for reference. **F)** Summary graphs of enriched neighborhood clusters, derived from their enriched N-Orbits, with corresponding example cell type maps. Solid edges indicate 1-orbit relationships between N-Orbits and dashed edges indicate 2-orbit relationships. Edge weights indicate relative frequency of cell type interactions among enriched N-Orbits. Self-edges indicate monotypic interactions of that cell type.

HDBSCAN clustering on the top 140 neighborhoods, as determined through feature selection and PCA loadings, revealed an NR-enriched cluster (p = 9.18e-5, 53 neighborhoods), an R1-enriched cluster (p = 8.63e-9, 44 neighborhoods), and an R1/R2-enriched cluster (p=0.011, 41 neighborhoods) that was most prominent at T3 (R1/R2-T3) (Figure 6E). The summary graph for the NR-enriched cluster showed clique formation among malignant epithelial cells, macrophages, and stromal cells, as well as interactions between malignant cells and various lymphocyte cell types including regulatory T cells (Figure 6F, Supplementary Figures 10-12). Although Shiao *et al*. did not report these interactions, prior research has indicated that interactions between malignant TNBC cells, tumor-associated macrophages (TAMs), and stromal cells such as CAFs contribute to tumor progression and recurrence in TNBC.^27^ In addition, CAFs have been shown to induce a lipid-associated macrophage phenotype that promotes immunosuppression in TNBC.^28^ Interestingly, the R1/R2-T3 cluster also contained the epithelial-macrophage-stromal cell clique observed in the NR cluster. However, unlike the NR cluster, regulatory T cells were absent from the R1/R2-T3 cluster, and the immune cells interacted only with the stromal cells and macrophages rather than directly with malignant epithelial cells. This suggests a potentially more effective anti-tumor mechanism that targets accomplice normal cells instead of malignant cells directly. The presence of plasma-epithelial cell interactions in the NR cluster aligns with findings from Wang *et al*. as well. Similar to the pCR summary graph from the Wang *et al*. dataset, plasma cells in the R1/R2-T3 cluster interacted only with normal cells, including accomplice cells. Lastly, the R1 cluster was characterized by interactions between B cells and CD4+ T cells, consistent with the formation of tertiary lymphoid structures, a distinguishing feature of this response group noted by Shiao *et al*.^29^ Additionally, macrophages in this cluster interacted in a monotypic manner, unlike their interactions with malignant cells in other clusters.

### Trajectory and homology reconstruction using N-Orbit distances

We utilized the N-Orbit distance metric to identify analogous brain regions across time in the aging mouse brain. The dataset from Allen *et al*. includes 31 MERFISH samples of the frontal cortex and striatum from 12 mice at 3 ages (4, 9, 24 weeks), with corresponding cell type and brain region annotations (Figure 7A-B).^30^ Annotated brain regions, provided by Allen *et al*., served as neighborhoods, totaling 241 across all samples. Using N-Orbit distances, we constructed a multipartite graph to visualize connections among neighborhoods across the three ages (Figure 7C). Our analysis revealed strong continuity of connections between neighborhoods representing the same brain region over time, with most cross-regional connections occurring within cortical and olfactory layers.

**Figure 7:**
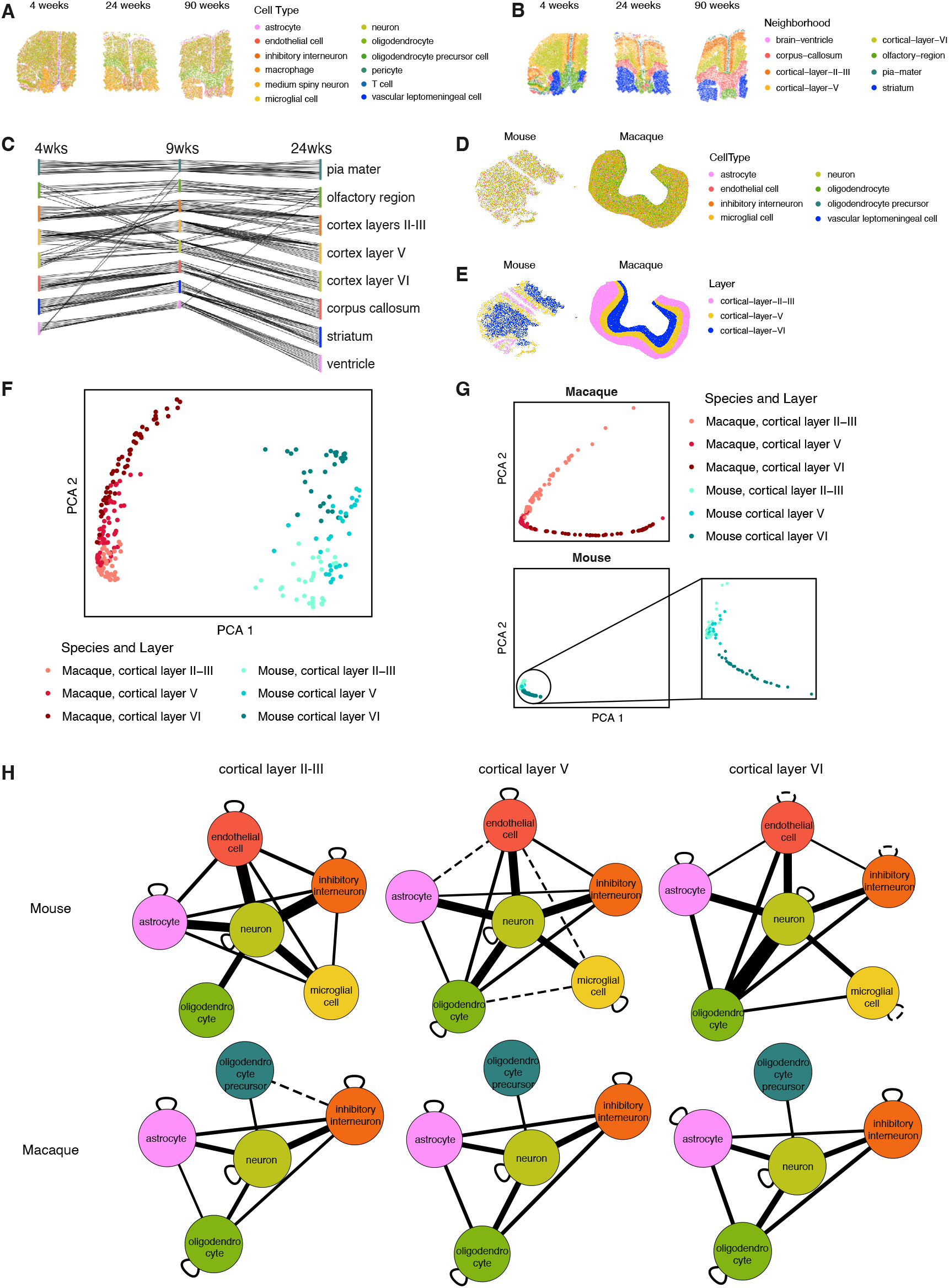
Application of N-Orbit distances to reconstruct mouse brain aging trajectories and mouse-macaque cortex homology. **A)** Cell type maps of example MERFISH mouse brain samples at each age time point. **B)** Neighborhood maps of example MERFISH mouse brain samples at each time point. **C)** Reconstructed trajectory graph of mouse brain region neighborhoods across ages, edges connecting each neighborhood node to the closest neighborhood (by N-Orbit-based neighborhood distance) in the adjacent time point. **D)** Cell type maps of example frontal cortex samples from mouse (MERFISH) and macaque (Stereoseq). **E)** Neighborhood maps of example frontal cortex samples from mouse (MERFISH) and macaque (Stereoseq). **F)** PCA plot of the N-Orbit-based neighborhood distance matrix for mouse and macaque cortex, color-coded by species and cortex layer. **G)** PCA plots of CTE-based neighborhood distance matrix from mouse and macaque cortices, color-coded by species and cortex layer, including a zoomed-in view of mouse cortical neighborhoods. **H)** Pruned summary graphs of each species and cortex layer, derived from enriched N-Orbits. Solid edges indicate 1-orbit relationships between N-Orbits, while dashed edges indicate 2-orbit relationships. Edge weights indicate relative frequency of cell type interactions among enriched N-Orbits. Self-edges indicate monotypic interactions of that cell type.

We further tested whether N-Orbit distances could capture homology between analogous regions across different species (Figure 7D-E). Using a macaque cortex dataset from Chen *et al*., in conjunction with the mouse brain dataset from Allen *et al*., we compared layered regions of the frontal cortex.^31^ The Chen *et al*. dataset includes 51 Stereo-seq samples across slices from one macaque brain. Cell type annotations in the macaque dataset were simplified to match those from the Allen *et al*. mouse brain dataset. Across both datasets, there were 246 neighborhoods annotated by their respective frontal cortex layers. We projected the neighborhood distance matrix for both datasets onto two dimensions using PCA (Figure 7F). The first principal component distinguished species (mouse vs. macaque), while the second principal component strongly corresponded to cortex layer, forming an ordered pattern across both species. This layer-specific pattern did not hold when using CTE-based distances (Figure 7G). Pruned summary graphs for each species and cortex layer revealed consistent cell type membership within each species, with variable cell type interactions across cortex layers (Figure 7H).

## DISCUSSION

### Summary of N-Orbit and advantages of approach

In this work, we presented a mathematical model and associated metric, N-Orbit, that enables principled comparison of TCNs. Unlike methods that rely solely on cell type composition and enrichment, the N-Orbit model can capture complex interactions between cell types within a TCN. In addition, the use of N-Orbits provides better practicality compared to the use of subgraphs, due to consideration of only proximity between cell types rather than the exact subgraph topology. This mathematical construct also allows for a vector representation of the N-Orbit structure, facilitating efficient algebraic manipulation and scalable computation of N-Orbit and neighborhood distances. Our results demonstrate that N-Orbit-based neighborhood distances effectively captured variations in spatial organization across various tissue types, outperforming cell type enrichment-based methods. Moreover, our findings suggest that the N-Orbit distance metric is broadly applicable across diverse biological contexts, including healthy, aging, and diseased organs, as well as homologous structures across different species.

### Impact of a neighborhood distance metric

The development of a neighborhood distance metric significantly broadens the scope of neighborhood analyses on spatial omics data. For neighborhood detection methods whose outputted labels are unaligned across samples, this metric enables clustering and identification of recurring neighborhoods across datasets. It also allows for the projection of neighborhoods onto a common coordinate space, allowing for the detection of patterns associated with temporal and clinical variables. In addition, the N-Orbit structure has high generalizability, as it only requires discrete units (e.g. cells) with spatial and phenotype information, focusing on proximity rather than precise connections between units. Consequently, N-Orbit has broad applicability in spatial analysis across diverse biological contexts, ranging from examining spatial organizational changes during embryological development to tracking the evolution of neighborhood-level structures within the tumor microenvironment. We also demonstrated the utility of N-Orbit distance-based approaches for predicting clinical outcomes using two triple-negative breast cancer datasets, each with distinct treatment approaches. Lastly, because our neighborhood distance metric is derived from a well-defined N-Orbit structure, neighborhoods and neighborhood clusters of interest can be traced back to their characteristic N-Orbits, allowing for interpretability of the results.

### Limitations

The design of the N-Orbit structure entails a few requirements. The input spatial omics data must be at single-cell resolution and include cell type annotations. Because the enumeration of N-Orbits is based upon neighborhood-filtered spatial k-Nearest Neighbors graphs, neighborhoods must exhibit a certain degree of spatial coherence to avoid overly sparse N-Orbit structures. Consequently, the N-Orbit model is more compatible with neighborhood detection methods that emphasize spatial coherence such as graph neural network-based methods like CytoCommunity and UTAG.

The performance of the N-Orbit neighborhood distance metric can vary based on the quality of results from the upstream neighborhood detection method. In our validation on the mouse spleen dataset, all six neighborhood detection methods demonstrated significant differences in distance between similar and different neighborhood types. However, classification performance as measured by AUROC showed considerable variation. Detection methods that produced partitions closely aligned with the ground truth, such as CytoCommunity and UTAG, achieved high AUROC scores (1 and 0.958, respectively). In contrast, other methods exhibited only moderate performance, with SpatialLDA yielding the lowest AUROC of 0.668. The quality of cell type annotations may also influence performance, though no cases were observed in this study where poor annotations significantly impacted the results.

The resolution of neighborhood distance matrices for projection, clustering, and prediction is influenced by sample size. Between the two TNBC datasets analyzed, the Wang *et al*. dataset had a much larger sample size (though smaller cell count per sample), resulting in nearly 15,000 total neighborhoods. In contrast, the Shiao *et al*. dataset included only 629 neighborhoods. This difference in sample size likely contributed to the difference in prediction performance, with the Wang *et al*. dataset achieving nearly perfect performance but not the Shiao *et al*. dataset. Despite the smaller sample size–comprising only 78 CODEX samples across 28 patients, including only 7 non-responders–the Shiao *et al*. dataset still achieved a macro-F1 score above 0.8.

## Acknowledgments

The authors thank the Children’s Hospital of Philadelphia Research Information Services for providing computing support. This work was supported by the National Institutes of Health (NIH) Human Biomolecular Atlas Program grant under award #U54 HL165442 (K.T.) and the National Cancer Institute (NCI) Human Tumor Atlas Network grant under award #U2C CA233285 (K.T.). KT holds the Richard and Sheila Sanford Endowed Chair at CHOP.

## Author Contributions

B.X. and K.T. conceived and designed the study. B.X. implemented the N-Orbit framework and wrote the software. B.X., Y.H. and K.T. performed data analysis. K.T. supervised the overall study. B.X. and K.T. wrote the manuscript with input from all authors.

## Supplemental Figure Legends

**Supplementary Figure 1:** Performance evaluation of N-Orbit model to distinguish synthetic neighborhoods of similar marginal but different joint distributions (Experiment 2).

**Supplementary Figure 2:** Hyperparameter and robustness testing for the N-Orbit model.

**Supplementary Figure 3:** Additional neighborhood distance UMAP density plots from the Wang *et al*. TNBC dataset.

**Supplementary Figure 4:** Full summary graphs for enriched neighborhood clusters from the Wang *et al*. TNBC dataset.

**Supplementary Figures 5:** List of enriched N-Orbits from the pCR-enriched neighborhood clusters from the Wang *et al*. TNBC dataset.

**Supplementary Figures 6:** List of enriched N-Orbits from the RD-B/O/P neighborhood clusters from the Wang *et al*. TNBC dataset.

**Supplementary Figures 7:** List of enriched N-Orbits from the RD-O/P neighborhood clusters from the Wang *et al*. TNBC dataset.

**Supplementary Figures 8:** List of enriched N-Orbits from the RD-P neighborhood clusters from the Wang *et al*. TNBC dataset.

**Supplementary Figure 9:** Additional neighborhood distance UMAP density plots from the Shiao *et al*. TNBC dataset.

**Supplementary Figures 10:** List of enriched N-Orbits from NR neighborhood clusters from the Shiao *et al*. TNBC dataset.

**Supplementary Figures 11:** List of enriched N-Orbits from R1 neighborhood clusters from the Shiao *et al*. TNBC dataset.

**Supplementary Figures 12:** List of enriched N-Orbits from R1/R2-T3 neighborhood clusters from the Shiao *et al*. TNBC dataset.

## Supplemental Table Legends

**Supplementary Table 1:** List of datasets analyzed in this manuscript.

**Supplementary Table 2:** List of p-values and ROC-AUC values from benchmarking experiments described in Figures 1-4.

**Supplementary Table 3:** List of parameters used for N-Orbit distance calculation.

## METHODS

### N-Orbit construction

We started with single-cell resolution spatial omics data, where each cell had assigned cell type and spatial coordinates. To decompose a neighborhood into its constituent N-Orbits, we first constructed a 6-Nearest Neighbors graph based on spatial coordinates, with nodes labeled by cell types and edges filtered to include only those within a 50μm distance. In the case where TCNs were the unit of interest, this graph was then filtered to nodes belonging to cells within each neighborhood. If samples were the unit of interest, filtering was skipped. For each cell in the graph, an *N*-hop structure was derived by identifying all cells within *N* edges from the center cell. N-hops were then converted to N-Orbits by considering the cell type of the center cell (*nucleus*) and the set of cell types within each n-hop, rather than the exact edges in the subgraph. For example, the set of cell types one edge away from the center cell formed the 1-orbit, while cell types two edges away, not included in the 1-orbit, formed the 2-orbit, and so forth. The depth of an N-Orbit, which indicates the number of orbits in the structure, was determined by the parameter *N*. Empty N-Orbits–those containing no cell types at any orbit level–are discarded.

### Vector representation of N-Orbits

We represented N-Orbits as vectors for computational analysis. This vector representation had two components: a nucleus encoding and an orbit encoding. Each component had an element for every possible cell type in the dataset, with indices arranged in alphabetical order. The nucleus encoding was set to zeros for all indices, except at the one corresponding to the center cell type, which was assigned a parameter value, *p*, described in the *Distance between N-Orbits* section. The orbit encoding assigned higher values for cell types that were closer to the nucleus. For instance, in N-Orbits with a depth of *N* = 2, cell types within the 1-orbit had a value of 2, while those within the 2-orbit had a value of 1. Cell types not present in the N-Orbit structure were assigned a value of 0.

### Distance between N-Orbits

The distance between a pair of N-Orbits was calculated as the Manhattan distance between their vector representations, corresponding to the number or cost of moves it would take to convert one N-Orbit into another by shifting cell types between orbits or outside of the N-Orbit structure. The nucleus change penalty parameter, *p*, determines the cost of changing the center cell type and can be adjusted to be higher than the cost of changing cell types within the orbits.

### Calculation of N-Orbit-based neighborhood and sample distance

The overall distance between two neighborhoods (or samples) was calculated as the minimum cost to convert the set of N-Orbits representing the first neighborhood (or sample) to the set of N-Orbits representing the second neighborhood (or sample). To ensure these sets are of the same size, we used a bootstrap of N-Orbits with size *s* (ranging between 1,000 to 20,000, depending on the average neighborhood or sample size) for each neighborhood (or sample). A cost matrix optimization using SciPy’s Linear Sum Assignment function was used to compute the minimum distance between these two sets of N-Orbit vectors.^32^ Let *D* be an *s* × *s* distance matrix between the N-Orbits representing the two neighborhoods. Let *X* be an *s* × *s* Boolean matrix representing a bijection between the N-Orbits of the two neighborhoods, where *X*_*ij*_ = 1 if the *i*-th N-Orbit in the first neighborhood was mapped to the *j*-th N-Orbit in the second neighborhood. The pairwise neighborhood distance was given by

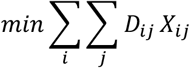

Distances for every possible pair of neighborhoods (or samples) can be compiled into an overall neighborhood (or sample) distance matrix. To account for N-Orbit sample size, distances were scaled by dividing by *s*.

### Calculation of cell type enrichment-based neighborhood and sample distances

We developed a similar schema for calculating neighborhood and sample distance based solely on cell type enrichment. The negative logarithms of p-values from hypergeometric testing for each possible cell type in a neighborhood (or sample) were compiled into a *cell type enrichment vector*. For p-values below the minimum value representable in Python (1e-309), the negative log p-values were recorded as 309. The distance between neighborhoods (or samples) was then calculated as the Manhattan distance between their respective cell type enrichment vectors.

### Synthetic neighborhood generation

Spatial coordinates for cells in the synthetic neighborhoods were generated using a Poisson process model (Figure 2A-E). Neighborhoods were assigned to cells by seeding one cell representing each neighborhood at random and allowing neighborhoods to expand along a 6-Nearest Neighbors spatial graph. The rate of spread was proportional to the pre-specified neighborhood composition within the sample. Following neighborhood assignment, cell type assignments were generated using a pairwise Markov random field.^33^ This method parameterizes the joint distribution of the labeled 6-Nearest Neighborhood graph based on node potentials ϕ(*x*_*i*_) and edge potentials ψ(*x*_*i*_, *x*_*j*_), where *x*_*i*_ denotes the cell type of node *i*. The joint distribution for *x* = {*x*_1_, …, *x*_*n*_} was given by

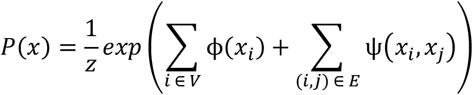

where *z* is a normalization constant that ensures the probability distribution sums to 1, and *V* and *E* are the sets of graph vertices and edges, respectively. In cases where cell types were independently distributed due to uniform edge potentials, the relative cell type proportions, or marginal distributions, were determined by the softmax of the node potentials

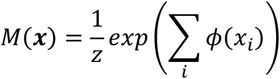

Cell type labels in the graph were initialized according to this marginal distribution. After initialization, cell types were resampled in random order according to the joint distribution.

To estimate the proper node potentials that would yield the target cell type composition *M*_*T*_ with the specified edge potential, node potentials were adjusted iteratively until the target parameters were approximated, as follows.

1. Node potentials were initialized as the logit transformation of the target marginal.
2. A sample was generated according to the specified edge potentials *ψ*(*x*_*i*_, *x*_*j*_) and current node potentials *ϕ*_*k*_(*x*_*i*_).
3. The empirical marginal distribution of the sample *M*_*e*_ was calculated along with its difference from the target marginal Δ*M* = *M*_*e*_ − *M*_*T*_.
4. The new node potential was formulated based on Δ*M* and a pre-specified learning rate α = 0.1.

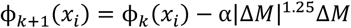
5. This process was repeated until either |Δ*M*| < 0.01 or 1000 iterations were completed.

Once the proper node and edge potentials were approximated for each neighborhood, these parameters were used to generate the specified number of synthetic samples. For each of the two simulated experiments, 50 samples–each consisting of roughly 10,000 cells–were generated for each sample type.

### Comparing distance metrics with AUROC

The ability of the N-Orbit and CTE distance metrics to distinguish between two groups (e.g. similar vs. dissimilar neighborhoods) was assessed based on their performance as single-variable classifiers, as measured by the area under the receiver operating characteristic curve (AUROC). The AUROC value was calculated from the true and false positive rates (TPR and FPR) at various thresholds, as follows:

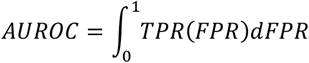

To compare the ROC curves and assess significant differences in performance between N-Orbit and CTE distance-based classifiers, we used the DeLong test (from *pROC* R package), along with the corresponding p-value. In some cases, the tests yielded p-values below the minimum representable number in R, which were recorded as p < 1e-324.

### Neighborhood instances in CODEX mouse spleen data

Neighborhoods in the mouse spleen CODEX dataset were divided into neighborhood *instances* by constructing a 25-Nearest Neighbors spatial graph with nodes labeled by neighborhood type. For each neighborhood, the graph was then filtered to include only nodes with the corresponding label, and each resulting connected component was designated as a separate neighborhood instance.

### Neighborhood instances in mouse hypothalamus MERFISH data

For each sample, an x-axis threshold was manually determined to split each sample into left and right halves. For all non-midline structures, cells within each hypothalamic nuclei region on either side of the threshold were designated as separate neighborhood instances.

### TNBC treatment response prediction models

For the TNBC treatment response prediction model on the Wang *et al*. dataset, a support vector machine (SVM) model with a radial basis function (RBF) kernel and L2 regularization was trained on rows of the neighborhood distance matrix reduced via PCA (# components = 35), along with treatment phase (Baseline, On-Treatment, Post-Treatment) and arm (chemotherapy only vs. chemotherapy + immunotherapy). Each row of the neighborhood distance matrix represented a vector of distances from one neighborhood to all other neighborhoods in the dataset. Fifteen-fold cross validation was used to assess test accuracy and record predictions for each neighborhood. Samples of underrepresented classes were given higher weight in the loss function such that each class had balanced weights in total. Patient-level predictions were made by ensembling neighborhood-level predictions associated with each patient and calculating the majority vote. In cases of a tie, the prediction accuracy contribution for the patient was set to 0.5. The prediction model for the Shiao *et al*. dataset followed the same framework except that it excluded the treatment arm variable and used 50 PCA components. CTE-based prediction models were constructed using the same method.

### Feature selection and neighborhood clustering

Recursive feature elimination (RFE), based on Scikit-Learn’s permutation_importance function, was used to rank the importance of PCA-derived features in predicting treatment outcomes from pre-treatment neighborhoods. The relative contribution of each original feature in the neighborhood distance matrix– each corresponding to a specific neighborhood–was calculated based on the most important features identified by RFE (3 for the Wang *et al*. dataset and 2 for the Shiao *et al*. dataset). The top contributing neighborhoods were then clustered using HDBSCAN clustering on their filtered neighborhood distance matrix. Neighborhood clusters enriched for different treatment outcomes were identified via hypergeometric test.

### N-Orbit enrichment test

The sampled N-Orbit vectors generated during the neighborhood distance calculation were used to identify enriched N-Orbits in the neighborhoods of interest, ensuring equal representation of all neighborhoods in the cluster. Enrichment was determined through a series of bootstrap-permutation tests. For each test, 20% of N-Orbit vectors were bootstrapped from the overall set and compiled into a matrix, with rows representing N-Orbit vectors. The nucleus encoding portions remained fixed, while the indices of the non-zero values within the orbit encoding portions were permuted. The indices corresponding to the 2-orbit (with a value of 1) were permuted first, followed by those of the 1-orbit (with a value of 2). If there were overlaps in resulting indices, the 1-orbit indices would override the 2-orbit indices. Remaining values not resulting from the permuted indices remained zero. This permutation procedure approximately preserved the marginal distribution of cell types and the distribution of N-Orbit cardinalities. For each unique N-Orbit in the original set, the occurrences in the unpermuted and permuted bootstraps were compared. Across 10 million tests, the proportion of tests where the count of the N-Orbits in the unpermuted bootstrap exceeded that in the permuted bootstrap was recorded as the p-value. Multiple testing correction was performed using the Benjamini-Hochberg method.

### Summary graph generation

Summary graphs were generated based on the set of enriched N-Orbits. A node was included for each cell type involved in any of the enriched N-Orbits. Solid edges were drawn for all 1-orbit relationships in the enriched set, weighted by the number of appearances across the N-Orbits. Dashed edges were drawn for all 2-orbit relationships that were not already captured by 1-orbit connections. Self-edges were drawn to indicate monotypic relationships of each cell type. *Pruned* summary graphs were generated by filtering to include nodes associated with edges of weight greater than 1 and nodes participating in cliques of size 3 or larger. All visualizations were created using Cytoscape.

